# Life and death of proteins after protease cleavage: protein degradation by the N-end rule pathway

**DOI:** 10.1101/115246

**Authors:** Nico Dissmeyer, Susana Rivas, Emmanuelle Graciet

**Author notes:** For correspondence; Twitter: @NDissmeyer.

## Abstract

The activity and abundance of proteins within a cell are controlled precisely to ensure the regulation of cellular and physiological processes. In eukaryotes, this can be achieved by targeting specific proteins for degradation by the ubiquitin-proteasome system. The N-end rule pathway, a subset of the ubiquitin-proteasome system, targets proteins for degradation depending on the identity of a protein N-terminal residue or its post-translational modifications. Here, we discuss the most recent findings on the diversity of N-end rule pathways. We also focus on recently found defensive functions of the N-end rule pathway in plants. We then discuss the current understanding of N-end rule substrate formation by protease cleavage. Finally, we review state-of-the-art proteomics techniques used for N-end rule substrate identification, and discuss their usefulness and limitations for the discovery of the molecular mechanisms underlying the roles of the N-end rule pathway in plants.

## I. Introduction: diversity of the N-end rule pathways

The control of protein stability plays a key role in the regulation of all cellular processes and, in eukaryotes, is largely controlled by the ubiquitin-proteasome system (UPS). The N-end rule pathway is a subset of the UPS and relates the *in vivo* half-life of a protein to the nature of its N-terminal amino acid residue and some of its post-translational modifications (PTMs). Removal of the initiator Met residue by methionine aminopeptidases (MetAPs) or cleavage of pre-proproteins (*i.e.* proteins that bear signal peptides and/or that require cleavage for their activation or degradation) by endoproteases exposes new N-terminal residues, potentially directing the resulting protein fragments for degradation by the N-end rule pathway (**Fig. 1**). Notably, the initiator Met residue may also serve as degradation signal for the N-end rule pathway (Hwang *et al*., 2010; Kim *et al*., 2014).

**Figure 1.**
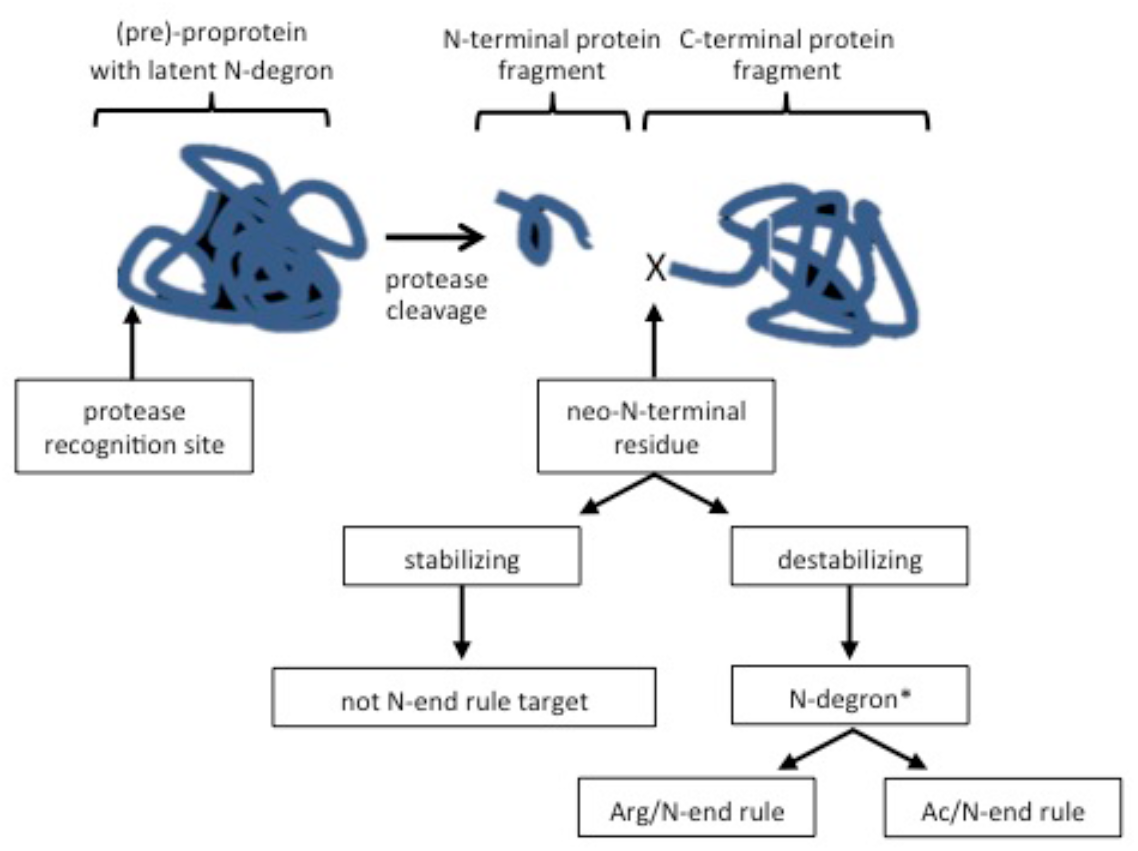
Role of proteases in the generation of N-end rule substrates. Endoproteolytic cleavage of a pre-proproprotein results in the exposure of a new N-terminal or ‘neo-N-terminal’ residue. This endoproteolytic event can expose to the solvent a ‘latent’ or ‘dormant’ N-degron that was previously buried in the internal sequence of the protease recognition site. Neo-N-terminal residues may be either ‘stabilizing’ or ‘destabilizing’ based on the Arg/ or Ac/ or Pro/N-end rule and may hence serve to target the protein fragment for degradation by the N-end rule pathway. Not all N-terminal destabilizing residues lead to degradation of the target protein. Additional structural and sequence features, such as flexibility of the N-terminal region, presence of Lys side chains as ubiquitin acceptor sites and charge or hydrophobicity of residues close to the neo-N-terminal are also critical for an N-end rule target.

In eukaryotes, the N-end rule pathway comprises different branches that mediate the degradation of proteins whose N-terminal residues are acetylated (Ac/N-end rule) and non-acetylated, respectively (reviewed in (Varshavsky, 2011; Gibbs *et al*., 2016)). Recent findings have revealed that the acetylation-independent branch of the N-end rule pathway was more complex than initially surmised. It now comprises two branches, the “classic” Arg/N-end rule (described in more detail below), and the newly found Pro/N-end rule pathway, which targets for degradation proteins with Pro at first or second position through the activity of the N-recognin Gid4, a subunit of the GID ubiquitin ligase (Santt *et al*., 2008) that targets gluconeogenic enzymes in yeast (Hammerle *et al*., 1998; Chen *et al*., 2017). While it is not yet known whether the newly found Pro/N-end rule is present in multicellular eukaryotes, components of the Arg/ and Ac/N-end rule pathways have been found to be mostly conserved within eukaryotes (reviewed in (Varshavsky, 2011; Tasaki *et al*., 2012; Lee *et al*., 2016)). For example, the complex hierarchical organization of the Arg/N-end rule pathway is similar in plants and animals (Graciet & Wellmer, 2010). N-terminal primary destabilizing residues can be directly bound by specific E3 ligases called N-recognins (termed PROTEOLYSIS1 (PRT1) and PRT6 in plants; **Fig. 2**). N-terminal secondary destabilizing residues require conjugation of Arg (a primary destabilizing residue) by the conserved Arg-transferases (ATEs). Lastly, N-terminal tertiary destabilizing residues are first enzymatically or chemically transformed into secondary destabilizing residues before arginylation by Arg-transferases (**Fig. 2**).

**Figure 2.**
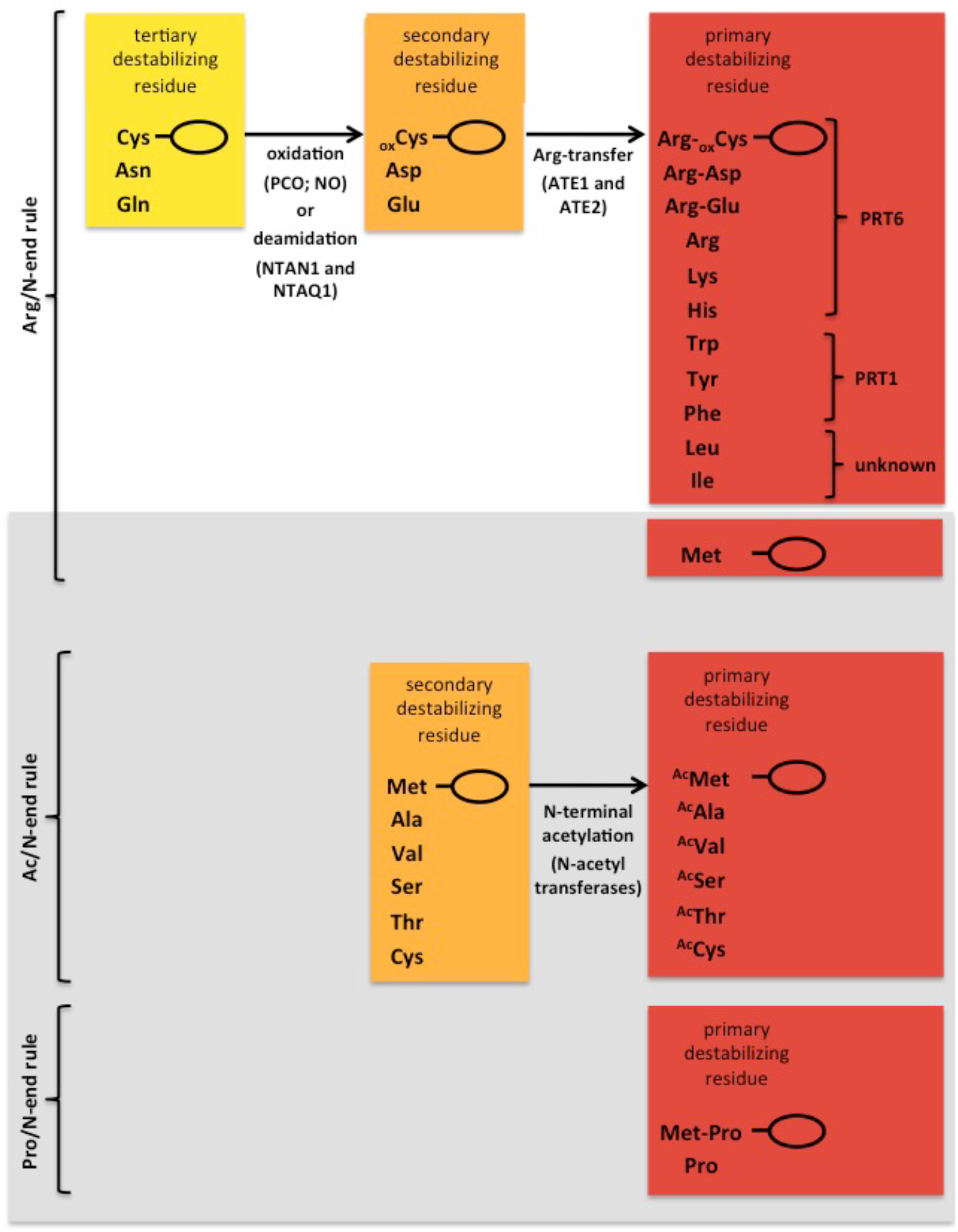
Canonical N-end rule pathways. The neo-N-terminus of a protein may be a tertiary (yellow), secondary (orange) or primary (red) destabilizing residue. One potential outcome of the exposure of neo-N-termini is their modification involving deamidation or Cys oxidation, arginylation, acetylation and finally ubiquitylation, followed by degradation of the protein by the Arg/ or Ac/N-end rule pathway. Protein fragments starting with primary destabilizing residues can be recognized and bound by so-called N-recognins, which belong to the class of E3 ubiquitin ligases. Note that the Ac/ and Pro/N-end rule (represented on a grey background) have not yet been formerly demonstrated to exist in plants, but are found in yeast (Hammerle *et al*., 1998; Chen *et al*., 2017). The Ac-N-end rule was also found in animals (Hwang *et al*., 2010; Varshavsky, 2011). Ovals denote the C-terminal protein fragments after (endo)proteloytic cleavage. NO: nitric oxide. *In vivo* studies with artificial N-end rule reporter substrates show that PRT6 in specific for positively charged residues (Garzon *et al.*, 2007) and that PRT1 is specific for aromatic hydrophobic residues (Potuschak *et al*., 1998). Furthermore, in *vitro* ubiquitylation assays of fluorescently labeled artificial N-end rule substrates confirmed the specificity of PRT1 and its E3 ligase activity (Mot *et al*., 2017).

The evolutionary conservation of the N-end rule pathway across different kingdoms of life is further underlined by the recent suggestion that chloroplasts (Rowland *et al*., 2015; Zhang *et al*., 2015) and mitochondria (Vogtle *et al*., 2009; Calvo *et al*., 2017) might also have specific N-end rule pathways that are more closely related to that of prokaryotes (see section III below).

## II. Defensive functions of the N-end rule pathway in plants

The Ac/ and the Arg/N-end rule pathway have been involved in the response to a variety of developmental and environmental signals (reviewed in (Gibbs *et al*., 2014; Gibbs *et al*., 2016; Lee *et al*., 2016)). A recent function identified in plants, is the PRT1-dependent degradation of the organ-size regulator BIG BROTHER, following cleavage by the protease DA1 in Arabidopsis (Dong *et al*., 2017). Interestingly, this work has uncovered the first substrate of PRT1.

Recent reports have described the involvement of the Ac/ and Arg/N-end rule pathways in the control of plant defense responses. First, the stability of the Nod-like immune receptors (NLRs) SUPPRESSOR OF NPR1, CONSTITUTIVE1 (SNC1) and RESISTANCE TO *P. syringae* pv. *maculicola1* (RPM1) is regulated through N-terminal acetylation (Xu *et al*., 2015). For example, NatA-mediated acetylation of the first Met residue of SNC1 contributes to its degradation. It is therefore tempting to speculate that the Ac/N-end rule pathway might participate to NLR homeostasis. Second, a new role for the Arg/N-end rule pathway was uncovered in activating the production of defense-related metabolites, such as glucosinolates and the phytohormone jasmonic acid (de Marchi *et al*., 2016). It was additionally shown that the Arg/N-end rule pathway positively regulates defenses against a wide range of bacterial and fungal pathogens with different lifestyles and, more particularly, that the Arg-transferases regulate the timing and amplitude of the defense program against avirulent bacteria (de Marchi *et al*., 2016). Third, a recent report highlighted a link between the known functions of the Arg/N-end rule pathway in the degradation of key transcriptional regulators of hypoxia response (the ERFVII transcription factors (Gibbs *et al*., 2016)) and *Arabidopsis* infection by the protist *Plasmodiophora brassicae,* which triggers clubroot development (Gravot *et al*., 2016). This study strongly suggests that Arg/N-end rule-driven hypoxia responses may be a general feature of pathogen-induced gall development in plants.

Altogether, these recently discovered defensive functions of the N-end rule pathways in plants are coherent with a potential role of defense-related plant proteases (including MetAPs) in generating N-end rule substrates. Indeed plant proteases are known to play important roles during plant/pathogen interactions (van der Hoorn & Jones, 2004). One potential example highlighting the connection between proteases and the generation of N-end rule substrates during plant/pathogen interactions is the cleavage of the central *Arabidopsis* defense regulator RPM1-INTERACTING PROTEIN4 (RIN4) by the *Pseudomonas syringae* protease effector AvrRpt2, which leads to RIN4 fragments with N-terminal destabilizing residues (Chisholm *et al*., 2005; Eschen-Lippold *et al*., 2016). Although no *in vivo* evidence has been provided to date, it has been suggested that AvrRpt2-derived RIN4 fragments are degraded by the Arg/N-end rule pathway (Takemoto & Jones, 2005).

Despite recent progress in our understanding of the Arg/N-end rule pathway and N-terminal acetylation in plants, the elucidation of the underlying molecular mechanisms has remained largely elusive. This delay is mostly due to the difficulties encountered to identify N-end rule substrates, in part because most N-end rule substrates are probably generated through protease cleavage of (pre-)proproteins.

## III. Proteases and degradation by the N-end rule pathway

Two recent proteomics studies highlight the prevalence of proteolytic events in plants. The first study (Zhang *et al*., 2015) used quantitative proteomics to identify N-end rule substrates that were expected to accumulate in Arg/N-end rule mutants compared to the wild type, while the second study (Venne *et al*., 2015) aimed at advancing techniques to characterize and quantify neo-N-termini for dissecting proteolytic events. Despite using different methods, these two studies showed that, in *Arabidopsis,* the majority of N-terminal fragments identified were the result of a protease cleavage or of initiator Met excision.

Notably, these two studies also found that most of the newly exposed N-terminal residues were not destabilizing based on the Arg/N-end rule, suggesting that (i) many proteolytic fragments are not substrates of the Arg/N-end rule pathway; (ii) fragments starting with destabilizing residues may not be detected, possibly because of their rapid Arg/N-end rule-dependent degradation. Furthermore, N-terminal acetylation accounted for 55% of the N-terminal fragments identified, with most of these appearing to be acetylated co-translationally. These results hence highlight the potential importance of the Ac/N-end rule pathway in plants (Zhang *et al*., 2015).

In agreement with the results mentioned above, most Arg/N-end rule substrates identified in yeast and animals are generated following protease cleavage (**Fig. 1 and 2)** (Rao *et al*., 2001; Ditzel *et al*., 2003; Piatkov *et al*., 2012a; Piatkov *et al*., 2012b; Brower *et al*., 2013); reviewed in (Tasaki *et al*., 2012)). However, in plants, our knowledge of protease cleavage sites and substrates is vastly lacking. Moreover, an N-terminal destabilizing residue is not necessarily sufficient for N-end rule mediated degradation, as suggested by the recent finding that many proteins with a presumed N-terminal destabilizing residue are relatively abundant and stable in plant cells (Li *et al*., 2017). The accessibility of the N-terminal residue for N-recognin binding, residues neighboring the N-terminus and the proximity of a Lys residue that may be ubiquitylated are also important (Tasaki *et al*., 2012; Wadas *et al.,* 2016).

The only putative N-end rule substrates that may be predicted using the primary sequence are proteins starting with Met-Cys, as MetAPs may excise the initial Met residue, exposing the Cys at the N-terminus of the protein. The resulting N-terminal Cys residue may then be oxidized through either a chemical reaction or the activity of Cys oxidases. The latter, termed PLANT CYSTEINE OXIDASEs (PCOs) have so far only been found in plants (Weits *et al*., 2014) and generate N-terminal Cys sulfinic acid (White *et al*., 2016). Screening through the literature can also reveal potential N-end rule substrates. For example, the identification of METACASPASE9 substrates has yielded a list of potential Arg/N-end rule substrates in *Arabidopsis* (Tsiatsiani *et al*., 2013). However, *in vivo* evidence that any of these fragments are degraded the Arg/N-end rule pathway is still lacking.

Finally, interesting novel developments include the potential existence of a chloroplast-specific N-end rule pathway (Nishimura & van Wijk, 2015). Indeed, recent studies show that following cleavage of the chloroplast transit peptide, destabilizing residues (of the prokaryotic N-end rule pathway) are under-represented in nuclear-encoded chloroplast proteins (Rowland *et al*., 2015; Zhang *et al*., 2015). Together with the recent discovery of a chloroplast ortholog of the bacterial ClpS N-recognin (Nishimura *et al*., 2013), these results suggest that stromal-processing peptidases may play a role in the generation of chloroplast N-end rule substrates (Rowland *et al*., 2015). Strikingly, a mitochondrion-specific N-end rule with similarities to the prokaryotic N-end rule could also exist (Vogtle *et al*., 2009; Calvo *et al*., 2017).

## IV. New proteomics approaches for the identification of N-end rule substrates

Workflows in shotgun proteomics were developed to compare global protein abundance in wild-type versus N-end rule mutants but provided only general peptide coverage with no information on N-terminal residues (**Fig. 3**) (Majovsky *et al*., 2014). This makes it difficult to draw any conclusions about potential N-end rule substrates.

**Figure 3.**
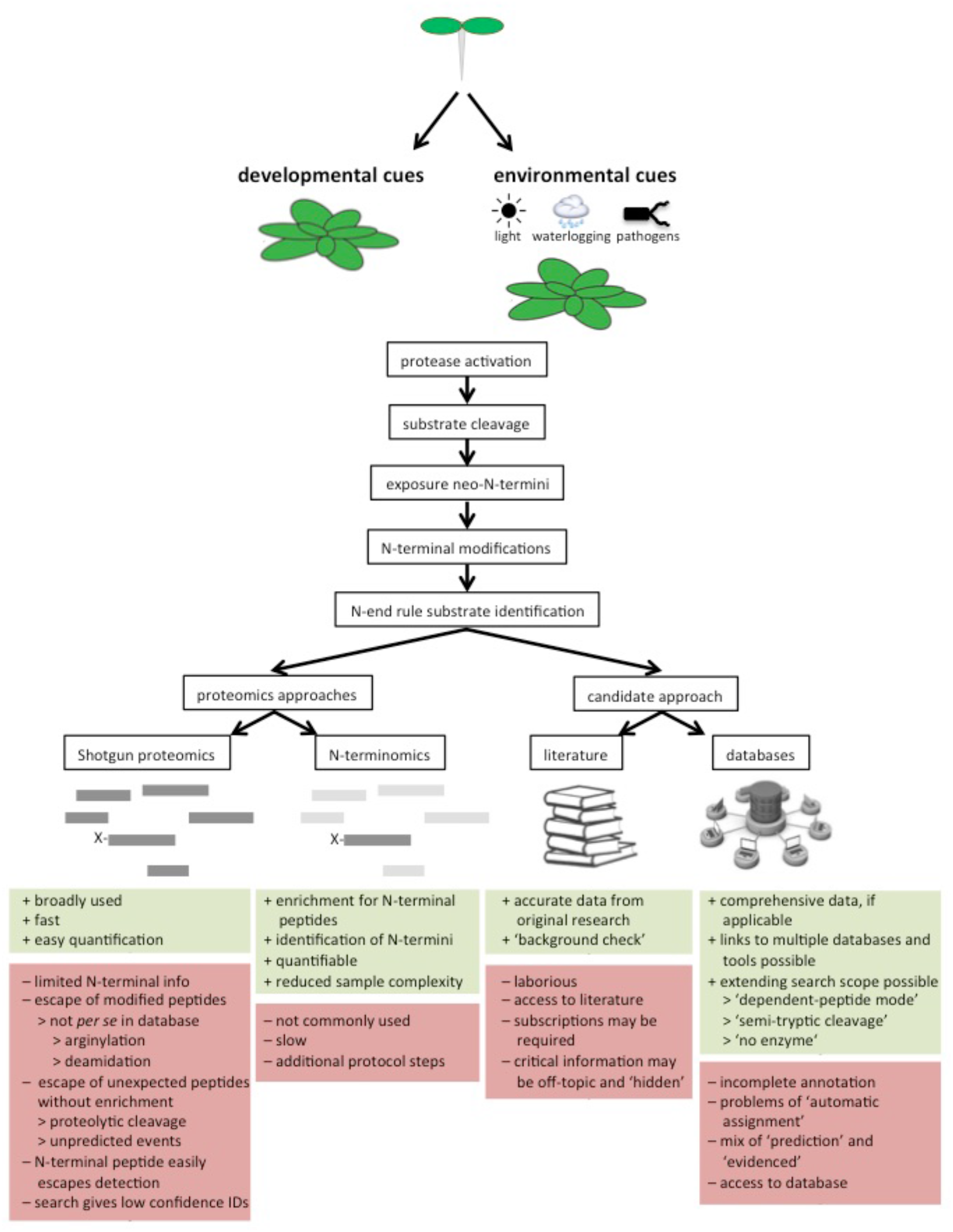
Approaches used to identify N-end rule substrate candidates. Endoproteolytic cleavage of (pre-)proproteins plays an important role in the generation of N-end rule substrates. The activity of proteases is tightly regulated and may depend on both developmental and environmental signals. Hence, specific N-end rule substrates may exist only in specific conditions or at specific developmental stages. As primary protein sequence information alone is not sufficient to predict N-end rule substrates, more complex proteomics methods have been recently applied. These include shotgun proteomics and the more specific N-terminomics approaches. In addition, scanning through the literature or searching through databases can lead to the identification of putative N-end rule substrates. Pros and cons of the different methods are highlighted in green and red, respectively.

More recently, specific proteomics techniques, termed N-terminomics, were developed to characterize proteolytic events and identify newly exposed neo-N-terminal residues and PTMs (acetylation, oxidation, deamidation, arginylation). These approaches are based on targeted enrichment of N-terminal peptides through chemical labeling of *α*-amine groups of N-terminal residues, which makes them distinguishable from internal amines derived from sample treatment by proteases (Huesgen & Overall, 2012). This specific N-terminal labeling reduces the complexity of the peptide mixtures and allows the identification of the true N-termini of mature proteins.

N-terminomics approaches use various strategies to separate N-terminal peptides from internal ones. COFRADIC (COmbined FRActional DIagonal Chromatography (Gevaert *et al*., 2003)) and ChaFRADIC (Charge-based FRActional DIagonal Chromatography (Venne *et al*., 2013)) use different chromatographic techniques to enrich for N-terminal peptides. TAILS (Terminal Amine Isotopic Labeling of Substrates (Kleifeld *et al*., 2010; Rowland *et al*., 2015; Zhang *et al*., 2015)) allows the capture N-terminal peptides via chemical modification. Other techniques include SILProNAQ (Stable-Isotope Protein N-terminal Acetylation Quantification (Bienvenut *et al*., 2015)) and PICS (Proteomic Identification of protease Cleavage Sites (Schilling *et al*., 2011)). As illustrated recently (Majovsky *et al*., 2014; Venne *et al*., 2015; Zhang *et al*., 2015), these techniques have the potential to discover candidate N-end rule substrates, which will greatly contribute to our understanding of the molecular mechanisms underlying the functions of the N-end rule pathway in plants (**Fig. 3**).

In another recent study, protein immunoprecipitation with an antibody specific for N-terminally arginylated sequences (i.e. modified by Arg-transferases) followed by mass spectrometry led to the identification of several candidate substrates of Arg-transferases in *Physcomitrella patens* (Hoernstein *et al*., 2016).

## V. Applications based on the N-end rule pathway

The major advances in our understanding of the N-end rule in plants have led to potential strategies for the agronomical improvement of crops. For example, it has been shown that mutants of Arg-transferases or PRT6 accumulated ERFVII transcription factors that act as master regulators of hypoxia response. This accumulation correlated with increased tolerance to waterlogging (Gibbs *et al*., 2011; Riber *et al*., 2015; Gibbs *et al*., 2016; Mendiondo *et al*., 2016).

In addition, the N-end rule pathway was used to conditionally target specific proteins for degradation using the temperature-labile nature of the low-temperature degron (lt-degron) (Faden *et al*., 2016). The lt-degron can be used as a small fusion tag to conditionally accumulate proteins of interest in living multicellular organisms including plants, so that their activity can be tuned by interfering with their degradation *in vivo.*

## VI. Concluding remarks

A key question in plants is to understand the roles of proteome dynamics, in particular how the regulation of protein stability contributes to developmental processes and to plant responses to environmental cues. In recent years, the N-end rule pathway has emerged as a key player in these processes. Genetic and biochemical approaches have revealed a wide range of functions in plants. Despite this major advance, which includes the discovery of the ERFVII transcription factors and BIG BROTHER as the first plant targets of this pathway, the number of potential N-end rule substrates identified remains small. This is largely the result of (i) technical limitations of proteomics approaches, (ii) the possibly restricted spatial and temporal accumulation of substrates, as the generation of these substrates often requires specific endogenous or exogenous triggers such as a stress or developmental cues that lead to endoproteolytic cleavage by proteases, and (iii) the lack of knowledge of protease cleavage sites in plants and the resulting neo-N-termini.

The potentially large field of N-end rule substrates generated by endoproteolytic cleavage is currently underexplored, partly due to the insufficient sensitivity of current proteomics methods and protocols. These bottlenecks, together with the nature and low abundance of PTMs to be detected (i.e. acetylation, oxidation, deamidation, arginylation) have hampered their discovery. Method optimization for the identification of N-end rule substrates includes the use of various proteases as alternatives to trypsin, which cleaves after Arg. Improving search algorithms that allow to identify all possible peptides, including unusual ones with mass increments corresponding to specific PTMs is also essential. Potential loss of information can be counteracted by using adapted search modes such as ‘semi-tryptic cleavage’ or ‘no enzyme’. These settings can better detect protein fragments which may result from unpredictable cleavages by yet unknown proteases and therefore presenting irrelevant aberrant peptides or false positives when using regular modes. Improved proteomics methods will in turn increase the completeness and accuracy of databases for protease cleavage sites, further facilitating the identification of N-end rule substrates.

Future improvements in proteomics will aim at analyzing N-terminal residues and their modifications by N-terminomics, thus contributing to the identification of N-end rule substrate candidates and hence to our understanding of the molecular mechanisms underlying N-end rule functions. For example, one function of the N-recognin PRT1 is related to the regulation of plant-pathogen responses, but the substrates responsible for the increased susceptibility of *prt1* mutants remain unknown (de Marchi *et al*., 2016).

In summary, the N-end rule pathway represents a central and emerging field of investigation to understand the role of protein degradation in plants. It has strong implications for our understanding of proteome dynamics and has an impact on traits relevant for agronomy (e.g. tolerance to waterlogging). Furthermore, easy manipulation of turn-over rates of recombinant target proteins by using temperature-inducible N-degrons indicates that the N-end rule pathway is also a valuable tool for biotechnology. We anticipate that the next developments in the field of system-wide N-terminomics will greatly contribute to the identification of N-end rule substrates.

## Acknowledgements

We thank Wolfgang Hoehenwarter (Proteomics Unit, Leibniz Institute of Plant Biochemistry, Halle), Ines Lassowskat (Institute of Biology and Biotechnology of Plants, University of M u nster), and Saskia Venne (Protein Dynamics, Leibniz Institute for Analytical Sciences, Dortmund) for helpful comments. Work in E.G. lab is funded by a Science Foundation Ireland award to EG (13/IA/1870) and through the Virtual Irish Centre for Crop Improvement (VICCI), which is funded by the Department of Agriculture, Food and the Marine (award 14/S/819). Work in N.D. lab is supported by a grant for setting up the junior research group of the *ScienceCampus Halle - Plant-based Bioeconomy* to N.D., by grant LSP-TP2-1 of the Research Focus Program “*Molecular Biosciences as a Motor for a Knowledge-Based Economy*’ from the European Regional Development Fund (EFRE) to N.D., by grant DI 1794/3-1 of the German Research Foundation (Deutsche Forschungsgemeinschaft, DFG) to N.D., by the Landesgraduiertenförderung Sachsen-Anhalt, the DFG Graduate Training Center GRK1026 *“Conformational Transitions in Macromolecular Interactions*'’ at Halle, and the Leibniz Institute of Plant Biochemistry (IPB) at Halle, Germany. Work at the LIPM is supported by the French Laboratory of Excellence project “TULIP” (ANR-10-LABX-41; ANR-11-IDEX-0002-02). E.G. and N.D. are participants of the European Cooperation in Science and Technology (COST) Action BM1307 – “European network to integrate research on intracellular proteolysis pathways in health and disease (PROTEOSTASIS)”.

